# Tolerance to NADH/NAD^+^ imbalance anticipates aging and anti-aging interventions

**DOI:** 10.1101/719401

**Authors:** Alvar J. Alonso-Lavin, Djordje Bajić, Juan F. Poyatos

## Abstract

Redox couples coordinate cellular function, but the consequences of their imbalances are unclear. This is somewhat associated with the limitations of their experimental quantification. Here we circumvent these difficulties by presenting a new approach that characterizes fitness-based tolerance profiles to redox couple imbalances using an *in silico* representation of metabolism. Focusing on the NADH/NAD^+^ redox couple in yeast, we demonstrate that reductive disequilibria generate metabolic syndromes comparable to those observed in cancer cells. The tolerance of yeast mutants to redox disequilibrium can also explain 30% of the variability in their experimentally measured chronological lifespan. Moreover, by predicting the significance of some metabolites to help stand imbalances, we correctly identify nutrients underlying mechanisms of pathology, lifespan-protecting molecules or caloric restriction mimetics. Tolerance to redox imbalances becomes thus a valid framework to recognize fundamental properties of the aging phenotype while providing a firm biological rationale to assess anti-aging interventions.

## Introduction

Research on redox homeostasis expanded substantially over the last two decades, continuously reshaping classical notions of oxidative cellular damage (Halliwell and Gutteridge, 2015). Among the most paradigmatic molecular agents underlying this homeostasis emerge the ratios of redox couples, like those of the conjugate forms of glutathione, NADPH and NADH. Both glutathione and NADPH act as essential scavenging mechanisms of reactive oxygen species (ROS) in mitochondria, while NADPH and NADH couple anabolic and catabolic pathways, respectively, with the redox state of the cell.

Even so, new mechanisms linking NADPH/NADP^+^ and NADH/NAD^+^ pairs to redox homeostasis continue to be recognized. For instance, the balance of NADPH/NADP^+^ partially explains the pro-survival consequences of AMP-activated protein kinase (AMPK) (She et al., 2014) and also associates circadian timekeeping with redox state (Rey et al., 2016). The NADH/NAD^+^ ratio is currently thought to be involved in the coordination of mitochondrial and nuclear function, the epigenetic regulation of DNA repair and cellular identity, and the tuning of energy metabolism to environmental variables (Cantó et al., 2015; Gomes et al., 2013). In non-pathological conditions, the NADH/NAD^+^ ratio fluctuates with environmental redox state, with hypoxic conditions and higher oxygen availability correspondingly co-occurring with reductive and oxidative deviations (Clanton, 2007; Graef et al., 1999).

But the growing interest in redox couple ratios mainly comes from their implications in pathology. The appearance of ROS in both the reductive (hypoxic, NADH prone) and the oxidative (hyperoxic, NAD^+^ prone) senses has been related to divergences from an optimal redox potential that ensures the best performance of the mitochondria (Aon et al., 2010; Clanton, 2007). Regarding cancer, decreased NADH/NAD^+^ may underlie lethality of glioblastomas (Gujar et al., 2016) and promote colon cancer progression (Hong et al., 2019), yet, it can also rescue some healthy phenotypes to different degrees in cells from other tumor types (Garrido and Djouder, 2017).

NADH has also become a point of interest in gerontobiology. In this context, the augmentation of the NAD^+^ pool resulted in the partial reversal of aging and otherwise trauma-related phenotypes across organisms (Das et al., 2018; Mendelsohn and Larrick, 2014; Wei et al., 2017; Zhu et al., 2017), and senescent and neoplastic cells have been found to present imbalances of the NADH/NAD^+^ ratio (Braidy et al., 2011; Schwartz and Passonneau; Wiley et al., 2016). Furthermore, the newly discovered roles of NADPH and the emergent concept of NADH/NAD^+^ as a master regulator of redox homeostasis and senescence are all in line with the metabolic stability theory of aging (Demetrius, 2004). This theory proposes that the cause of aging is the vulnerability of the steady-state levels of redox couples to random environmental perturbations on enzyme reaction rates, and makes several interesting predictions that apply to humans.

Given all these implications, many studies have examined the phenomenology of redox couple ratios either by passively reporting their levels or by actively modifying them. Experimental manipulations are however challenging. The most traditional accuse deep experimental caveats (Sun et al., 2012) and newer ones still miss on certain biological circumstances due to being restricted to temperature and pH intervals (Hung et al., 2011; Zhao et al., 2015). Moreover, it is experimentally expensive to monitor the wide array of phenotypes following alteration of coenzyme pools by metabolite supplementation (Hou et al., 2010) and mutations or over-expressions of NAD(H)-consuming enzymes (Bai et al., 2011; Felipe et al., 1998). Thus, there is a need for alternative strategies to address the control of redox homeostasis through the manipulation of redox couples, as well as our understanding of the biological consequences of this control.

*In silico* models become a practical research strategy whenever experimental approaches are limited, with the advantage of enabling a full mechanistic account of the observed phenomena. Genome-scale metabolic models, which can be studied through flux balance analysis (FBA)(Orth et al., 2010) have become a standard in systems biology for studying the consequences of metabolic perturbations on cellular function. Among other contributions, they have contributed to the discovery of new antibiotics and chemotherapeutics, the design of bacterial strains optimized for industrial production of substances of interest, and the better comprehension of human metabolic diseases (Burgard et al., 2003; Pagliarini et al., 2016; Raman et al., 2009). The use of flux balance analysis has the additional advantage of providing insight into metabolic phenomena without influence of non-metabolic confounding factors (genetic, epigenetic, mechanical, etc.). Thus, genome-scale metabolic models are particularly well suited to examine the metabolic consequences of deviations from redox homeostasis.

Here, we use FBA to probe the balance of redox couples on a genome-scale reconstruction of the unicellular eukaryote *Saccharomyces cerevisiae*, whereby we characterize the metabolic and longevity-related consequences of a controlled perturbation of the available NADH/NAD^+^ flux across different genetic backgrounds. More specifically, our results reveal that tolerance to this imbalance leads to a specific metabolic rerouting reminiscent of pathology and also explains more than a quarter of the intra-specific variability in post-mitotic lifespan. In addition, this framework helps us outline a computational protocol (that we also apply to animal and human metabolic models) for identifying metabolites and enzymes with potential as therapeutic targets in the context of age-related pathologies.

## Results

### A fitness-based tolerance profile characterizes redox couple perturbations

To represent an imbalance between the conjugate forms of a redox couple, we incorporated an artificial reversible reaction –the “imbalance reaction”– into the genome-scale reconstruction of the corresponding metabolic network (Methods). The reaction oxidizes or reduces the couple while considering specific cellular compartments (e.g. cytosol, mitochondria, etc.) and its activity can be fixed to any desired rate value. For any of these values, one can calculate a growth rate (“fitness”), which acts as a proxy for the tolerance of the yeast cell to that particular condition. Finally, a tolerance profile is defined by computing growth rate for a range of imbalance values (**Fig.1A;** note that reductive/oxidative conditions are represented in blue/red, respectively, throughout the manuscript).

**Figure 1.**
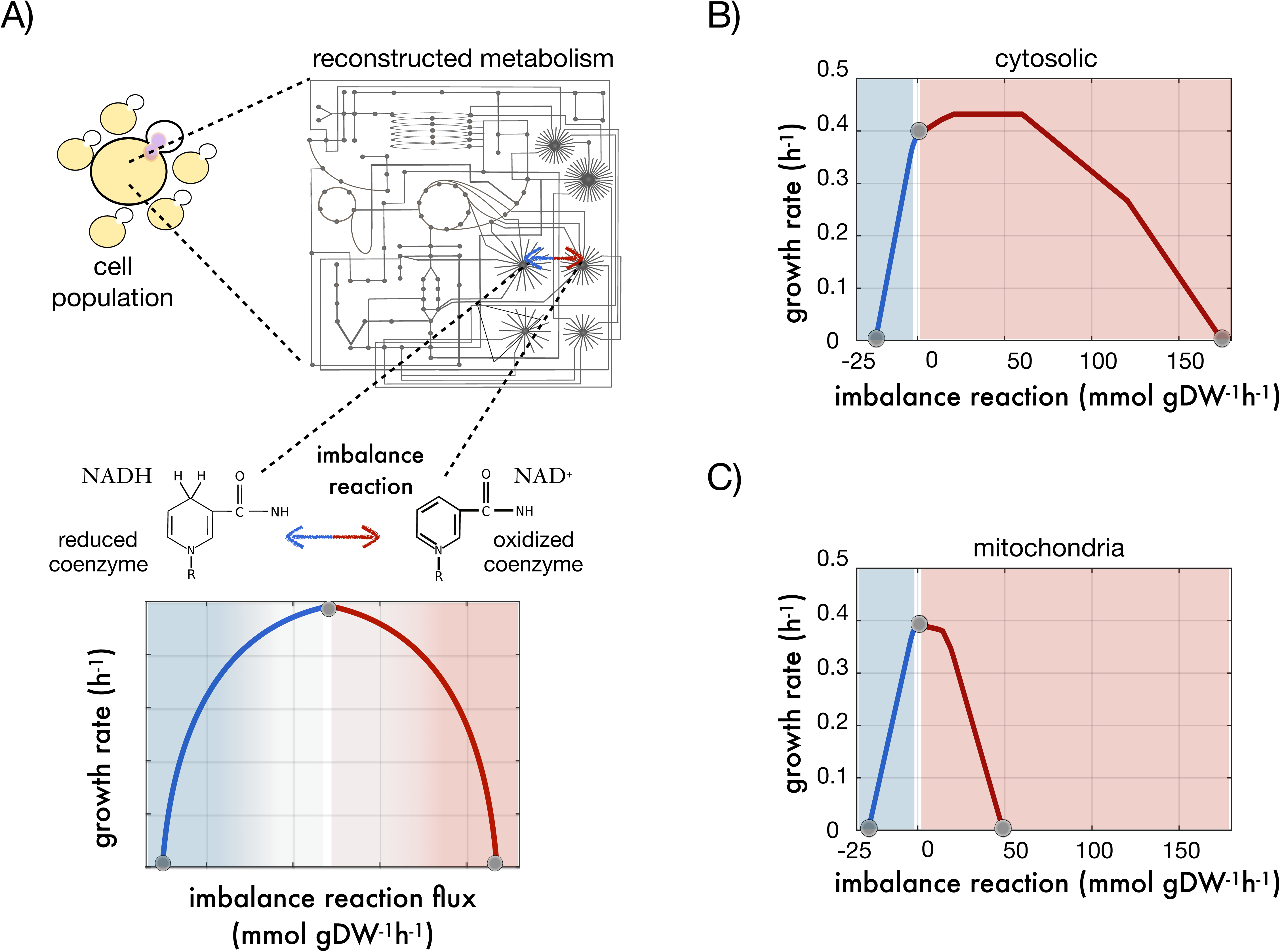
A tolerance profile characterizes the response to redox imbalances. A) Top. We introduced an artificial reaction in the metabolic reconstruction model of an organism (in this case yeast) to interconvert the two conjugate forms of a redox coenzyme (here the NADH/NAD^+^ pair). Under each of a series of imbalance conditions, i.e., rate values of the artificial reaction, we apply flux balance analysis to compute growth rate. Bottom. The predicted values of growth are plotted against the rate values of imbalance what delineates the tolerance profile; a proxy for the tolerance of the metabolism when faced with the chosen perturbation. B) Tolerance profile in yeast associated to imbalances located in the cytosol. C) Tolerance profile in yeast associated to imbalances located in the mitochondria. Blue/red shading represents reduced and oxidized imbalance regimes, respectively, and grey dots indicates values corresponding to no imbalance or extreme reductive/oxidative imbalances producing no growth.

Tolerance profiles typically exhibit a maximum growth around the null imbalance point, with roughly any deviation (i.e. nonzero value of the reaction) leading to decreased fitness. This emphasizes the fact that, for metabolism to work, the activity of the reactions that tune a redox couple ratio in one sense must be proportionate to the activity of those that tune it in the other. More specifically, cytosolic imbalances of NADH/NAD^+^ in *S. cerevisiae* growing on glucose and aerobic conditions yield a profile with a maximum growth significantly displaced towards the oxidative side on a point of the imbalance profile where ∼50 mmols/gDW/hour of NADH are being converted to NAD^+^ (**Fig. 1B**). When considering instead the imbalance in the mitochondria, we observed a maximum at the point of null imbalance (**Fig. 1C**), a pattern that we similarly observed in other profiles (**Fig. S1**). In general, reductive conditions become deleterious and lethal faster than oxidative regimes. In two cases, (the conjugated pairs of cytosolic NADH or mitochondrial Thioredoxin) mild artificial oxidation of the couple improves growth **Fig. S1**).

### NADH/NAD^+^ perturbations cause metabolic syndromes that are reminiscent of pathology

The energy metabolism of yeast at no imbalance corresponds to a characteristic aerobic metabolism in the presence of glucose (the growing conditions studied) in which glycolysis is coupled with the Tricarboxylic Acid Cycle (TCA) and oxidative phosphorylation. The pentose phosphate pathway oxidizes glucose and provides ribose-5P for nucleotide synthesis and NADPH-born reductive power for anabolism, while anaplerotic routes departing the TCA cycle, like those of glutamine metabolism, are moderately used to primarily feed pyrimidine and amino acid synthesis. FBA enables us to quantify changes in these pathways and how they eventually detail the metabolic traits underlying any particular imbalance regime.

Specifically, **Fig. 2A** shows how reductive imbalances of cytosolic NADH/NAD^+^ produced an increase in glycolytic flux, a decrease in the activity of the TCA cycle and the electron transport chain, and a rise of glutamine metabolism. This pseudohypoxic metabolic signature –in the presence of oxygen– resembles anaerobic metabolism, where glycolysis is coupled with alcoholic or lactic fermentation in detriment of mitochondrial pathways, the oxygenic part of the pentose phosphate pathway is shut down and glutamine metabolism, more active, might be rerouted to produce pyruvate on top of contributing to anabolism. Notably, this phenotype captures some features of paradoxical yield metabolisms observed in different types of cancerous cells (the Warburg effect) (Potter et al., 2016).

**Figure 2.**
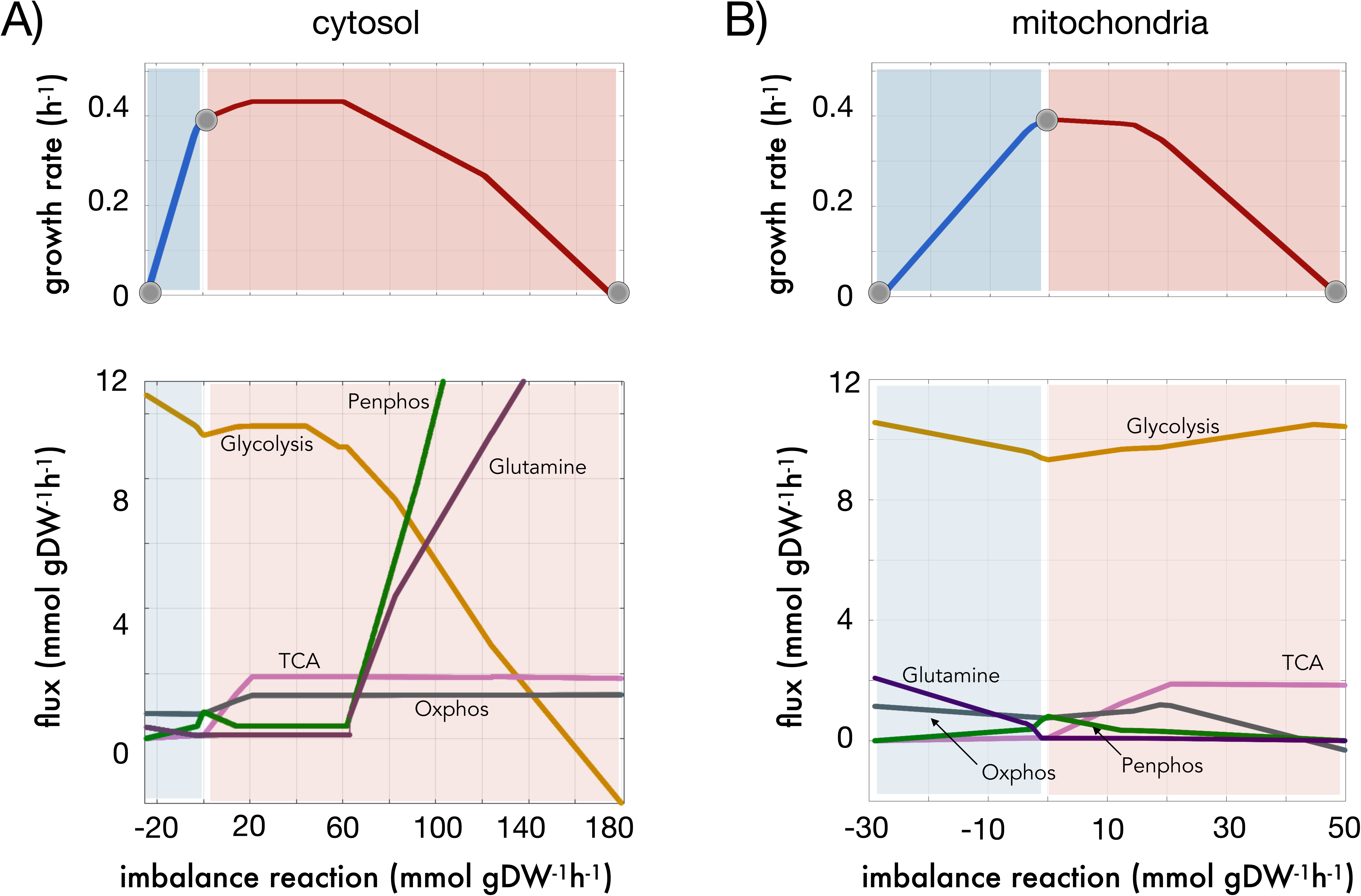
Fluxes of the main pathways of energy management underlying the tolerance profile in yeast. A) Cytosolic NADH/NAD^+^ imbalance (top) and flux values of five representative pathways (bottom): i/glycolysis (Glycolysis, ocre), ii/Krebs cycle (TCA, pink), iii/pentose phosphate (Penphos, green), iv/oxidative phosphorylation (Oxphos, gray), and glutamine metabolism (Glutamine, purple). The represented flux vectors are the result of averaging the flux of all reactions of the particular pathway. B) Same as A) with respect to mitochondrial imbalance. Note the presence of negative fluxes in glycolysis (panel A, bottom) represent increased gluconeogenesis. See main text for details.

In contrast, the energy metabolism underlying oxidative tolerance (with respect to cytosol, **Fig. 2A**) exhibited a more aerobic-like configuration but with peculiarities, such as a particularly active polyamine metabolism; and extreme properties, including increased gluconeogenesis, oxidative phosphorylation and TCA cycle activity, as well as a very high (up to 12 fold the normal level) flux through the pentose phosphate pathway.

When the imbalance reaction is located in the mitochondria, the reduction of NAD^+^ produced again a certain pseudohypoxic behavior, with one difference (**Fig. 2B**). The flux through glycolysis and glutamine metabolism increased, with a concomitant loss on the parts of the TCA cycle and the pentose phosphate pathway. However, and unlike in the cytosolic case, oxidative phosphorylation increased significantly. On the other hand, the oxidative side of the mitochondrial profile was more idiosyncratic: glycolytic activity increased in parallel with that of the TCA cycle, but oxidative phosphorylation worked for the most part at lower levels than normal, and glutamine metabolism was of little importance.

### Metabolic syndromes result from a compromise between redox balance, biomass production and an ATP/NADH trade-off

We identified several key elements at play that shaped the previous syndromes. The oxidative perturbation was met with an exacerbated aerobic response as a compromise between maintaining growth and buffering the imbalance perturbation. This involved rerouting flux through the maximum possible number of reactions that reduced NAD^+^ while preserving a global flux distribution that was capable of generating biomass constituents. These two mechanistic elements (perturbation buffering and biomass maximization) are the most relevant requirements of the optimization problem and sufficient to describe the oxidative regime of the tolerance profile.

The reductive side required however one additional insight. As more and more NAD^+^ is sequestered to NADH, reactions that use NAD^+^ and are directly or indirectly necessary to produce biomass constituents become more and more constrained, so energy metabolism must be rerouted to allow for an elevated conversion of NADH to NAD^+^ and to limit reduction of NAD^+^ to NADH. This is still insufficient to face the perturbation, as most reductive power in the form of NADH is essentially useless to many metabolic objectives, reactions and growth: the energy stored in NAD^+^ must be reallocated to ADP. Thus, metabolism must prioritize reaction modules that produce as much ATP and as little NADH as possible; it must rely on shunts and pathways that have a high ATP/NADH yield, e.g., glycolysis and oxidative phosphorylation.

This results, among other things, in reduced TCA cycle and increased glycolytic flux (**Fig. 3A**). To further explore the impact of this ATP/NADH trade-off, we overlapped a reductive NADH/NAD^+^ perturbation with an artificial reaction that allows phosphorylation of ADP. The simulations showed that the elevated glycolysis to TCA cycle flux ratio that characterizes reductive metabolism is dependent on the ATP/NADH yield (**Fig. 3B**). Forcefully phosphorylating ADP reduces this pseudohypoxic signature even in the face of very strong NADH-prone imbalance rates.

**Figure 3.**
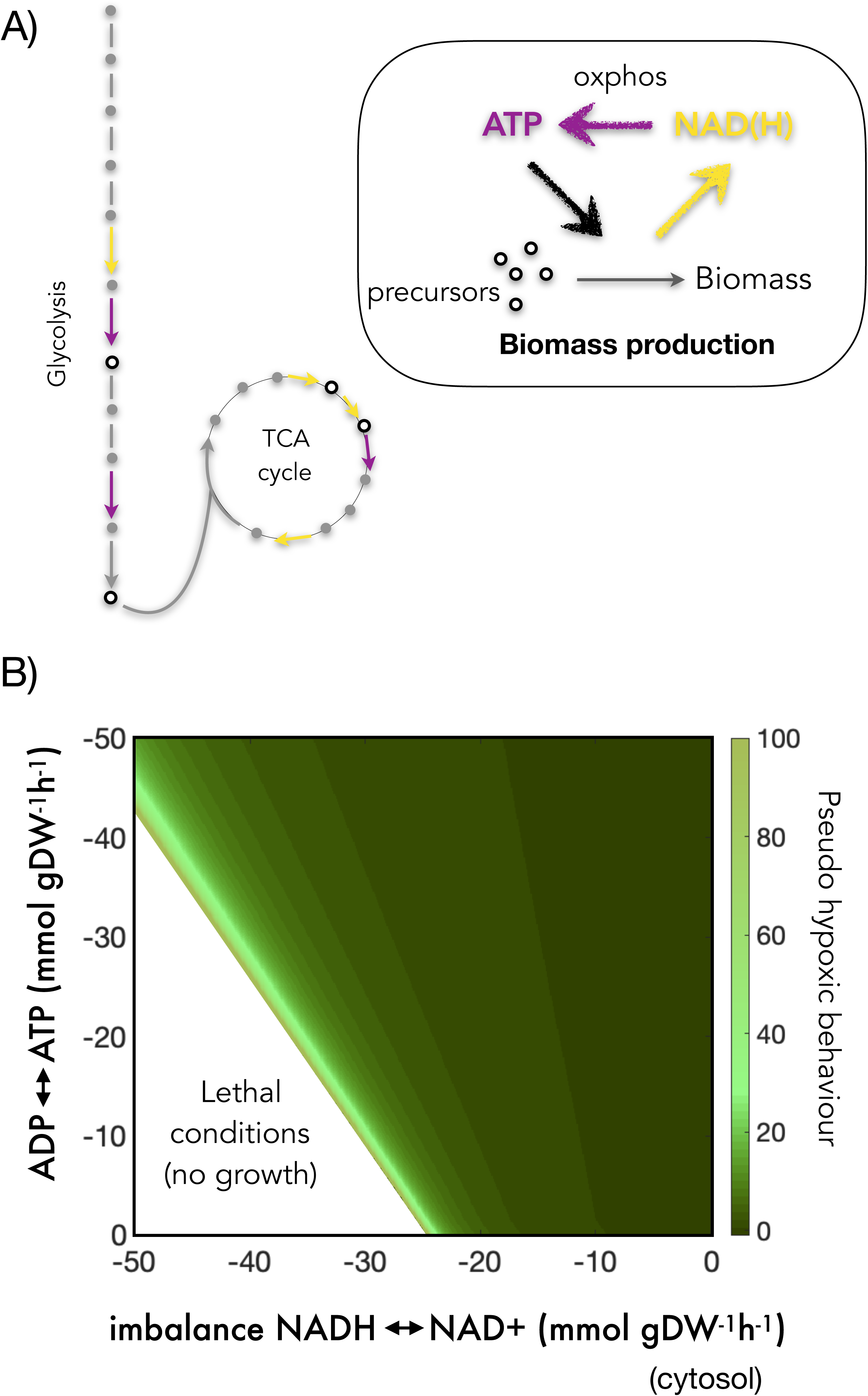
Competing mechanisms cause the pseudohypoxic behavior of yeast. A) Balance among NADH, ATP and biomass precursor production favors reaction modules that produce as much ATP and as little NADH as possible to compensate the consequences of reductive regimes, e.g., the use of glycolysis to that of TCA. Note here that purple arrows represent ATP production, yellow arrows represent NAD(H) production and white circles indicate generation of biomass precursors, B) An NADH-prone NADH/NAD^+^ perturbation (*x* axis) is overlapped with an artificial ADP phosphorylation reaction (*y* axis) that forcefully introduces reductive power in the form of ATP into the imbalanced metabolism. A green color gradient represents the ratio between glycolytic and Krebs cycle flux normalized by its normal value (up to 100-fold). It can be appreciated that ADP phosphorylation reduces the pseudohypoxic phenotype and delays quiescence.

### Tolerance explains experimental chronological lifespan differences between different yeast mutants

We asked next to what extent could the tolerance profile act as predictor of lifespan, given that redox couples have been discussed as potential lifespan determinants. One way to study this is to compute the profile in different mutants (**Fig. 4A**) and then quantify how it corresponds to exact lifespan measures, normalized chronological lifespans (CLS), available from experimentally measured mutant survival curves (Garay et al., 2014). CLS are calculated from these mutant survival curves as the increase in stationary phase survival relative to the wild-type.

**Figure 4.**
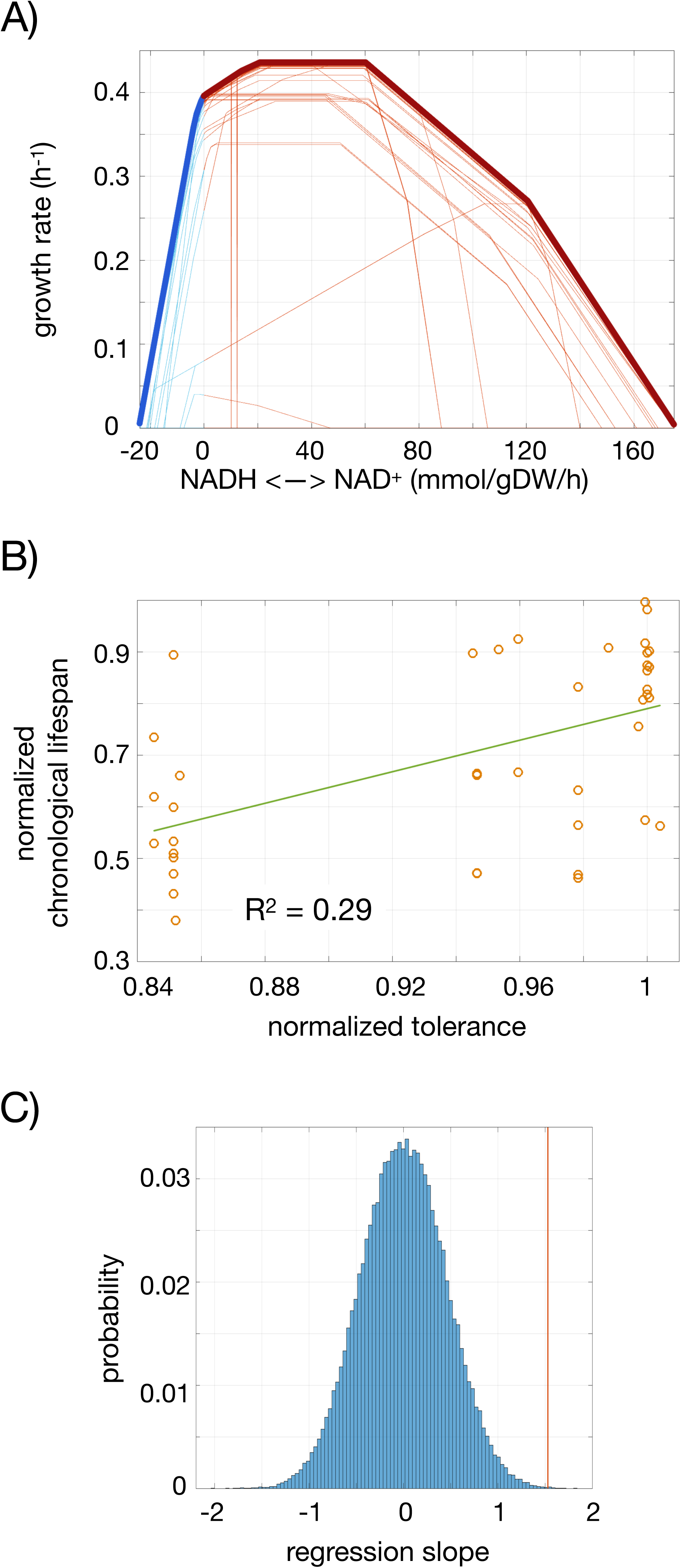
Tolerance score as predictor of chronological lifespan in yeast. A). Tolerance profiles obtained for yeast mutants; blue/red sectors of the curve represent the reductive/oxidative regime of the NADH/NAD^+^ disequilibrium. B) Association between normalized tolerance score (proportional to the breadth of the imbalance values at both oxidative and reductive regimes, Methods) and chronological lifespan. The correlation explains ∼30% of the total variance (R^2^ = 0.29, p-value = 3.2×10^-4^, *N* = 41). C) As an alternative way of view this association, we obtained a histogram of regression slope values obtained from ten thousand randomly generated associations between tolerance score and lifespan. From this sample, we find only 3 cases in which the association between the tolerance score and the lifespan data is stronger than the one found (indicated by the red vertical line).

In flux balance analysis, mutations in specific genes are simulated by constraining the flux of the reactions associated to them through Boolean rules that relate each chemical reaction to ORFs that translate for the reaction’s enzyme (Methods). For each of these mutants, we computed a mutant tolerance profile (**Fig. 4A**), and used the sum of the imbalance absolute values at which growth rate is halved (both in the reductive and the oxidative regimes) as a scalar score of tolerance.

Our mutant set was limited however by some restrictions (Methods). Notably, we were incapable of distinguishing differences in tolerance below 10 ppm of the wild-type’s value without reaching prohibitive computation times, and many mutants presented both negligible differences in lifespan and negligible differences in tolerance. Additionally, it is generally considered that FBA is incapable of characterizing gain of function deletions and, quite predictably, no mutant tolerances exceeded that of the wild-type.

Beyond these constraints, the *in silico* tolerance profiles were able to explain ∼30% of the experimentally measured lifespan variability (**Fig. 4B**, R^2^ = 0.29, *N* = 41, p-value = 3.2×10^-4^) with great significance: 10000 randomizations of the data pairs led to only 3 instances with a larger regression slope (**Fig. 4C**).

### Conventional nutrients enable tolerance to NADH/NAD^+^ imbalances

Lastly, we investigated whether specific dietary metabolites were particularly determinant in the response to redox imbalance. To this aim, we used an additional feature of FBA models, which is the possibility of accessing the usage of a particular metabolite (defined as the rate of consumption in the steady state, Methods). We examined the way this rate changed with increasing values of reductive and oxidative NADH/NAD^+^ imbalance.

Usage was rather linear in both sides of the profile and for most metabolites. Thus, we fitted this change pattern to a linear model and considered the (absolute) slope as a scalar representative of the corresponding metabolite’s relevance for tolerating redox imbalance (**Fig. 5A**). Among the top responsive nutrients of iAZ900 we noticed dietary metabolites that are known to play a critical role in regulating yeast lifespan, such as acetate (Burtner et al., 2009), as well as many that increase lifespan experimentally in yeast, worms or even human cells (Madeo et al., 2018; Mishur et al., 2016) including malate, hydroxybutyrate, spermidine or oxaloacetate (**Figs. 5B-D, Supplementary Table 1**).

**Figure 5.**
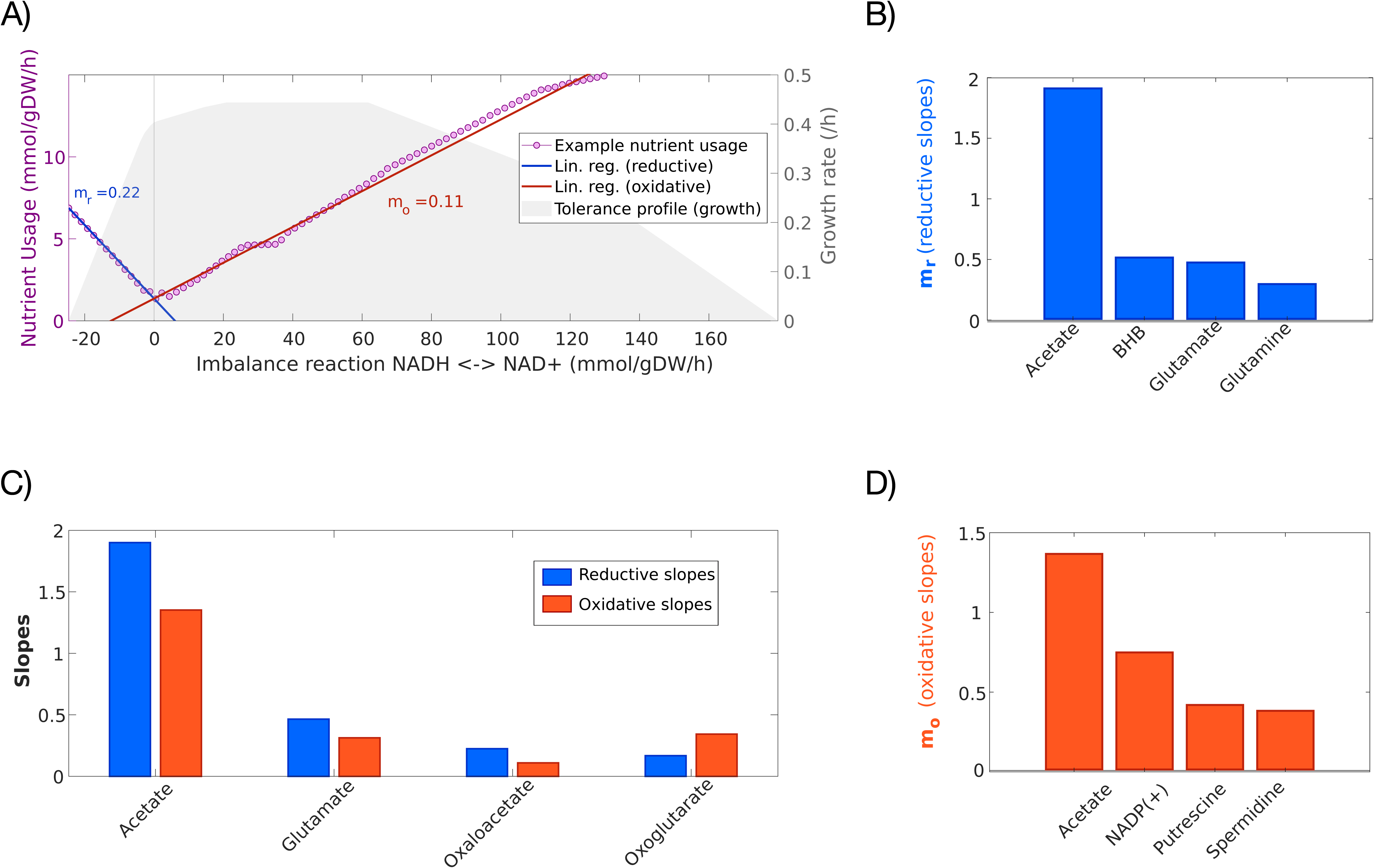
Homeostatic nutrients in yeast. A) Example usage profile of a nutrient with its corresponding reductive (blue) and oxidative (red) linear regressions characterized by slopes m_r_ and m_o_, respectively (in absolute values). We included the cytosolic tolerance profile in the background as a reference. B) Linear regression slopes (m_r_) of the top 4 homeostatic nutrients in the reductive sense of the NADH/NAD^+^ perturbation. C) Linear regression slopes of the top 4 nutrients that are homeostatic in both senses of the perturbation; Blue: Reductive linear regression slope; Red: Oxidative linear regression slope. D) Linear regression slopes (m_o_) of the top 4 homeostatic nutrients in the oxidative sense of the NADH/NAD+ perturbation.

Certain nutrients were more relevant for tolerance to NAD^+^ reduction, others to NADH oxidation, and a few to both of these regimes. The top most important dietary metabolites for reductive tolerance were in order acetate, beta-hydroxybutyrate (BHB), glutamate and glutamine (**Fig. 5B**), meanwhile the top most important for tolerating NADH oxidation were acetate, NADP^+^, putrescine and spermidine (**Fig.5D**). Among those that participated from tolerance on both sides of the profile, the most relevant were in order acetate, glutamate, oxaloacetate and oxoglutarate (**Fig.5C**).

We considered metabolic models in other organisms to further corroborate which nutrients are determinant in the response to imbalances. All these top contributors changed, albeit not widely, with alpha-keto acids, redox couples, certain vitamins and certain amino acids being significantly necessary to control NADH/NAD^+^ perturbations in *C. elegans* and the human reconstruction. The most prevalent reaction to redox imbalance in these organisms pertains metabolites that mediate pH homeostasis, such as acetate, bicarbonate, biphosphate, sodium, water and others of the like. Similarly, the relevance of glutamate, glutamine, aspartate, threonine, serine and glycine distinguishes them from other amino acids and most metabolites. Medium sized, oxidized acids like oxoglutarate, malate and oxaloacetate also play a role in tolerance consistently, as tend to do biotin and some folates (see **Supplementary Table 1** for a complete list).

## Discussion

We propose here an alternative approach to understand the broad biological consequences of changes in redox couples. This approach is based on *in silico* metabolic models and introduces the notion of the tolerance profile as a measure that quantifies the cellular resilience to these changes.

The metabolic adjustments underlying the profile reveal the presence of a pseudohypoxic phenotype associated to the reductive NADH regimes. This phenotype is reminiscent of some apparently paradoxical low yield energy metabolisms observed in cancers (the Warburg effect), and also recognized in yeast (Crabtree effect) and bacterial (overflow metabolism) cells (Basan et al., 2015; Mori et al., 2016; Potter et al., 2016). The possibility that this behavior could be caused by resource allocation constraints arising at comparatively high growth or glucose uptake rates has been put forward in recent years (Basan et al., 2015; Mori et al., 2016). However, the pseudohypoxic phenotype we observe is independent of growth rate and glucose uptake, and in fact, it co-occurs with low growth rates (Methods). We have shown that its cause lies on a fundamental ATP/NADH trade-off, a rationale that is supported by a recent experimental study (Maldonado and Lemasters, 2014).

Furthermore, our flux analysis suggests that ATP maintenance can be negatively affected by reductive NADH/NAD^+^ disequilibria. Increased NADH is thought to be a correlate of decreased ATP availability, as impairment of oxidative phosphorylation might result in both an increase in NADH/NAD^+^ and a decrease in ATP/ADP. We show that extrinsically generated NADH imbalances can be a cause of decreased energy availability through orthogonal metabolic mechanisms, even while oxidative phosphorylation works over normal levels. This is very significant in the context of aging research, as decreased energy availability and ATP/ADP ratios are a conserved hallmark of cellular aging and age-related pathologies (Moreira et al., 2003; Pall, 1990; Yaniv et al., 2013) and might promote the accumulation of toxic waste and the loss of proteostasis (another hallmark of aging) by decreasing protein turnover and therefore increasing protein half-life (Anisimova et al., 2018).

We aim next to determine the validity of our framework as a predictor of lifespan and of dietary metabolites buffering the redox imbalances. Tolerance does anticipate cellular lifespan, with some limitations due to the available dataset. Controlling for these limitations (Garay et al., 2014), we find that the resulting correlations are still enough evidence of a relationship between the variances of tolerance and CLS.

Contrary to our expectations, the most distinct lesson drawn from our analysis of dietary metabolites is that the main substances driving the response to the imbalance does not particularly rely on the NAD^+^ salvage network. Indeed, the top “homeostatic nutrients” are intermediates of the TCA cycle and other parts of central metabolism whose action is far more permeating than that of NAD^+^ precursors. In addition, the relevance of reactions that reduced or oxidized NAD(H) while acting as bridges between the redox pair and major metabolic pathways is far superior to that of NAD^+^ salvage limiting enzymes (such as nicotinamide mononucleotide adenyltransferase).

For instance, oxaloacetate and oxoglutarate score in the yeast model among the top four most effective metabolites underlying tolerance in both reductive and oxidative conditions, a consistent feature that corroborates prior experimental results (Chin et al., 2014; Williams et al., 2009). Other significant metabolites include hydroxybutyrate that has been consistently shown to increase lifespan, regulate NAD^+^ and mediate the response to starvation (Edwards et al., 2014; Newman and Verdin, 2014), and spermidine, which belongs to the family of polyamines and is known to play roles in age-related processes, autophagy and DNA protection (Eisenberg et al., 2009; Minois et al., 2011; Pietrocola et al., 2015).

We used *C. elegans* and human models to strengthen the previous evaluation, revealing a broader picture that is centered around pH homeostasis, redox couples, and the TCA cycle. This suggests that the ways in which pH (Burtner et al., 2009) and NADH imbalance (Ayer et al., 2014) determine senescence in cells are deeply intertwined. Beyond pH, the most pervasive and important nutrients to regulate NADH/NAD^+^ imbalance are the alpha-keto acids oxaloacetate and oxoglutarate, their aminated forms, and other mitochondrial-related metabolites like malate, pyruvate and fumarate, i.e. the main hub of redox balance control is the TCA cycle.

To this day, the mechanisms through which amino acids and TCA cycle intermediates impact life extension in yeast and *C. elegans* remain obscure. Metabolites like malate, oxaloacetate, fumarate, valine, serine or threonine can indeed increase the lifespan of organisms, but the processes leading to these effects are debated and complex (Edwards et al., 2015, 2013). Our results indicate that a common explanation to all these pro-longevity phenomena lies in the effect of the nutrients on the capacity of cells to tolerate perturbations in the NADH/NAD^+^ ratio.

One could argue however that some of the metabolites considered appear self-evident as they are after all involved in reactions that inter-convert NADH and NAD^+^. The question is then why other metabolites that also look *a priori* self-evident do not emerge in our results. The answer lies in the mechanisms that ensure realistic predictions in FBA. For a nutrient to be “homeostatic” against redox imbalance it must not only increase NADH or NAD^+^ production, but stand in central a pathway or module with a high ATP/NADH yield and/or capacity to provide biomass constituents.

Finally, two more insights from our results are noteworthy. On one hand, they suggest that in response to redox imbalances, metabolic networks are poised to increasingly produce and/or consume some metabolites that are interpreted by signaling networks as precluding the need for autophagy, antioxidant and hormetic responses; as well as many which excess or supplementation has been found to increase lifespan and/or otherwise mimic the effects of caloric restriction (CR), in a manner dependent on the signaling pathways that are implicated in CR-mediated lifespan extension. This reinforces previous evidence linking CR and NADH/NAD^+^ balance as part of the same lifespan-extending and health promoting process (Lin et al., 2004).

On the other hand, our study shows that in response to the altered ratios, metabolism also makes increasing use of certain substances that can chemically damage the cell, such as acetate, putrescine or acetaldehyde; as well as some that can promote tumorigenesis through metabolic rewiring, such as glutamine, succinate and fumarate (Sciacovelli et al., 2016). This could then partly explain the pathologies linked to redox imbalance and the macroscopic processes it is involved in, such as degenerative and oncologic diseases: if redox imbalance must be buffered with substances that are toxic, then these substances are probably mechanisms of the pathologies that co-occur with redox imbalance.

We realize that our approach to redox imbalance can be understood as an unusual variation of the study of metabolic network robustness and that it may accuse certain caveats that leave plenty of room for improvement. With respect to robustness, studies using FBA traditionally defined it as a change of the objective solution (typically growth) in response to varying reductions in the reaction rates, e.g. (Edwards and Palsson, 2000), rather than to a particular perturbation (redox imbalance) in the metabolites as we do. With respect to the limitations of our analysis, they can be linked to intrinsic limitations of FBA itself, like the absence of regulatory genes. Ultimately, the reliability of our results depends on the predictive power of the metabolic reconstructions: the current yeast models are predictive and advanced, but they are not perfect (Heavner and Price, 2015), and still they are far better than even the most accurate multicellular reconstructions available. Despite all these concerns, there is plenty of evidence that warrants the increasing fidelity of metabolic models to natural behavior.

Presently, the prevailing research tends to ignore the potential negative consequences of indiscriminately decreasing the NADH/NAD^+^ ratio. This is in part due to the promising benefits resulting from the mild decrements achieved experimentally, which include reduction of neoplastic phenotypes, lifespan, and healthspan extension. However, there is emerging evidence that recommends extreme caution regarding these positive results (Gujar et al., 2016; Hong et al., 2019), as well as a solid, experimentally sustained theoretical framework that predicts negative consequences from decreasing NADH/NAD^+^ ratios beyond a threshold (Aon et al., 2010). Our NADH/NAD^+^ imbalance tolerance profiles cater to this emerging picture, as mild oxidative deviations can be beneficial, but higher ones are as deleterious as the opposite extreme.

More specifically, our tolerance profiles suggest that on top of causing chemical or physiological problems, both low and high NADH/NAD^+^ ratios must also be met with purely metabolic drawbacks, including decreased energy availability and/or biosynthetic output. Furthermore, and as we have pointed out, the limited experimental observations that are available on some of the questions we tackle seem reminiscent of the results we report here.

## Acknowledgements

This work was supported by grant FIS2016-78781-R and the Salvador de Madariaga program (grant PRX18/00439) from the Spanish Ministerio de Economía y Competitividad (JFP).

## Authors contributions

A. A.-L., D.B. and J.F.P. designed the research, A. A.-L. performed the *in silico* experiments, A. A.-L. and J.F.P. analyzed the results and wrote the draft manuscript. All authors discussed and edited the final manuscript.

## Supplementary Figures and Tables

**Figure S1: Tolerance profiles of different coenzymes in yeast**. A) NADH/NAD^+^, cytosolic perturbation. B) NADH/NAD^+^, mitochondrial perturbation. C) NADPH/NADP^+^ cytosolic perturbation. D) NADPH/NADP^+^, mitochondrial perturbation. E) Thioredoxin, cytosolic perturbation. F) Thioredoxin, mitochondrial perturbation.

**Figure S2. Metabolic signature of redox imbalance in different organisms**. A) Pathway flux under NADH/NAD+ cytosolic imbalance in the *C. elegans* model iCEL1273. B) Pathway flux under NADH/NAD+ mitochondrial imbalance in the *C. elegans* model iCEL1273. C) Pathway flux under NADH/NAD+ cytosolic imbalance in the human metabolic reconstruction 2.02. D) Pathway flux under NADH/NAD+ mitochondrial imbalance in the human metabolic reconstruction 2.02.

**Table S1. Homeostatic nutrients**. Full list of ‘homeostatic’ nutrients, together with the strength of their effect (absolute value of slope) and the reliability of the linear fit (pearson’s R squared) for *S. cerevisiae, C. elegans*, and human metabolic models.

## STAR* Methods

### KEY RESOURCES TABLE

|  |  |  |
| --- | --- | --- |
| Software and Algorithms. |  |  |
| Matlab | Mathworks | https://www.mathworks.com |
| Cobra Toolbox | (Heirendt et al., 2019) | https://opencobra.github.io/cobratoolbox/stable/ |
| Gurobi | Gurobi Optimization, LLC. | http://www.gurobi.com |

### CONTACT FOR REAGENT AND RESOURCE SHARING

Further information and requests for resources, included computer code used in this study, should be directed to and will be fulfilled by the lead contact, Juan F. Poyatos.

### METHOD DETAILS

#### Metabolic models

We considered *S. cerevisiae* iAZ900 *in silico* genome-scale model as the basis for our study (Zomorrodi and Maranas, 2010). This model incorporates all the necessary complexity of *S. cerevisiae*’s metabolism (e.g., it is fully compartmentalized), has been empirically corroborated, and also reduces the computational load. Most calculations, like the tolerance profiles and metabolic signatures, were additionally tested on other models and species, including yeast iND750 (Duarte, 2004), *C. elegans* iCEL1273 (Yilmaz and Walhout, 2016) and the human metabolic reconstruction 2 (Swainston et al., 2016). We used the COBRA Toolbox (Heirendt et al., 2019) and the the Gurobi linear programming optimizer (http:/www.gurobi.com). Of note, some of the calculations are computationally expensive, so they need to be parallelized.

#### Imbalance implementation and tolerance profiles

We added an artificial reversible reaction to the models that specifically interconverts NADH and NAD^+^ (in a specific compartment) to represent an imbalance. Before each fitness calculation, we assign a rate value to the imbalance reaction, ensuring that in the steady state solution a certain quantity of flux of the reactions that produce one of the two forms is always lost to the conjugate form by virtue of its rerouting to the imbalance. By assigning precise values to the imbalance reaction, from a negative to a positive limit, we obtain the *tolerance profiles*. Note that negative values mean in this context that the reaction is working in reverse. Therefore, for a range of imbalance values, a fitness or growth vector (tolerance profile), and a reaction flux matrix (metabolic signature) are produced. The full extent of the calculations was carried out using only two imbalance reactions: a cytosolic and a mitochondrial one, but we also explored other compartments and tolerance profiles associated to different coenzymes (**Fig. S1**) (Thioredoxin, NADPH, and reactions placed in the nucleus, the extracellular medium or the endoplasmic reticulum) in each of the models we tested. Note also that when an imbalance reaction of a particular redox couple is placed in a compartment, it restricts the flux of each conjugate form that would otherwise be available for the chemical reactions of metabolism. This will impact reactions that use the redox couple in that same compartment directly. Reactions that use the redox couple in other compartments will be affected through changes in the transport rates of the conjugate forms.

#### Metabolic signatures and flux analysis

We grouped reactions into a particular subset of pathways: glycolysis, gluconeogenesis, the pentose phosphate pathway, oxidative phosphorylation, the tricarboxylic acid cycle (TCA), glutamine and glutamate metabolism, quinone biosynthesis, NAD^+^ salvage, and fermentation, alcohol and sterol metabolism. These routes described the basic catabolic and anabolic behavior of metabolism, but we noted peculiar reactions outside these subsystems when their flux was outstanding. To get a proxy for the total flux across a pathway we simply used the averaged flux of all reactions in the pathway. In each case, we also quantified variability to confirm that the mean is a representative score –with standard deviations being greatest in glycolysis and the pentose phosphate pathway. To quantify pseudohypoxia in **Fig. 3**, we adopted the ratio between the calculated fluxes of glycolysis and the TCA cycle, normalized to its wild-type (non-imbalanced) value.

#### Gene deletion study and tolerance score

For each gene available in the model, we performed a full effect (“homozygote”) mutation, removing all related enzymes from the model; then we ran one simulation for each value of the imbalance vector thus generating a *mutant* tolerance profile for each gene. From the resulting collection of mutant tolerance profiles, we eliminated those that were lethal in all conditions. We summarized the tolerance profiles into scalar scores by summing the imbalance absolute values (both oxidative and reductive regimes independently) at which the yeast’s maximum steady-state growth is halved with respect to the wild-type. This score was the one considered in the correlation with the yeast mutant chronological lifespan data from (Garay et al., 2014). For once, the effect on long termed viability of lethal mutations is by definition inaccessible. The experimental protocol for measuring yeast CLS is inaccurate for mutants that are affected in their rate of exponential growth (Garay et al., 2014), which makes data for such mutants unreliable. Some mutants have negligible differences both in lifespan and tolerance; to get a more accurate measure of the latter would require unrealistic computation times. Lastly, flux balance analysis cannot show gain-of-function phenotypes from loss-of-function mutations, and in this way cannot ascertain the tolerance increase of some mutants, which it represents as neutral. After controlling for all these groups, the resulting correlations are still enough evidence of a relationship between the variances of tolerance and CLS.

#### CAFBA study

To examine energy allocation constraints Constrained Allocation Flux Balance Analysis (CAFBA) was used upon the yeast models iND750 and iAZ900 following (Mori et al., 2016). The value of the ribosomal weight was obtained from the literature (Metzl-Raz et al., 2017), and the invariant sector was inferred from (Chagoyen and Poyatos, 2018) using a representative sample of 834 yeast proteins. We set the weights of the enzymes to the median value of several simulations and adjusted the weight of the proteome allocated to carbon uptake to correspond to a reasonably realistic glucose uptake rate of 15.5 mmol/gDW/h (Orth et al., 2010). The pseudo hypoxic phenotype generated by the imbalance affects different reactions in a different manner, e.g., CAFBA-elicited overflow is largely dependent on hexokinase flux, yet the overflow linked to the imbalance is quite resilient to changes in this enzyme’s activity.

#### Nutrient variation study

We calculated the total amount of metabolite that the steady state metabolism consumed *per* unit time for every imbalance situation, and then selected only the subset of nutrients whose usage increased with increasing values of either reductive or oxidative imbalance. Since all dependence profiles were reasonably linear and regime changes were tame and infrequent, we performed a linear regression on each side (reductive, oxidative) of each dependence profile and used the slopes of these regressions to be our proxy for how much each nutrient became important as redox imbalance increased. We then ordered the nutrients according to these slopes into three groups: nutrients important for tolerance in the reductive side of the profile, another for those that underlie oxidative tolerance, and a third group for the few nutrients that were used to counteract both reductive and oxidative imbalance. See

## TOLERANCE PROFILES ACROSS ORGANISMS ARE EXPLAINED BY SIMILAR METABOLIC RESPONSES

A comparison with other organisms can help evaluate the generality of the previous patterns (Methods, **Fig. S2**). The *Caenorhabditis elegans* metabolic response was very similar to that of yeast when the NADH/NAD^+^ imbalance reaction was placed in the mitochondria. The preceding pseudohypoxic behavior followed tightly that found in the yeast model, and so did most of the oxidative response. The only difference was that during very extreme oxidative imbalances the worm model drew heavily on a partial reverse configuration of the Krebs cycle (sometimes referred to as the glyoxylate cycle) (**Figs. S2A-B**).

The response of the worm model to cytosolic imbalances diverged slightly more from that of *S. cerevisiae*. For once, glutamine metabolism did not change much from its normal level under any point of the tolerance profile. Some features however did emerge in a similar fashion, like the increased glycolysis to Krebs cycle ratio in response to NAD^+^ reduction, and the elevated gluconeogenesis, pentose phosphate and oxidative phosphorylation flux following NADH oxidation (**Figs. S2A-B**).

In the case of the more intricate human metabolic network, we obtained a much wider tolerance profile, which made for more elaborate metabolic strategies and a more diverse response (**Figs. S2C-D**). However, some tendencies emerged that recover many of the patterns already described. Yet, the human recon differs from the other models in that these changes are far tamer, with fluxes rising and declining within a 10% to 2 fold of the normal reaction rates (compared to the much broader changes in *S. cerevisiae* and *C. elegans*).

