## Supplementary figures and images for "Tolerance to NADH/NAD^+^ imbalance anticipates aging and anti-aging interventions"

### Supplemental Figure 1

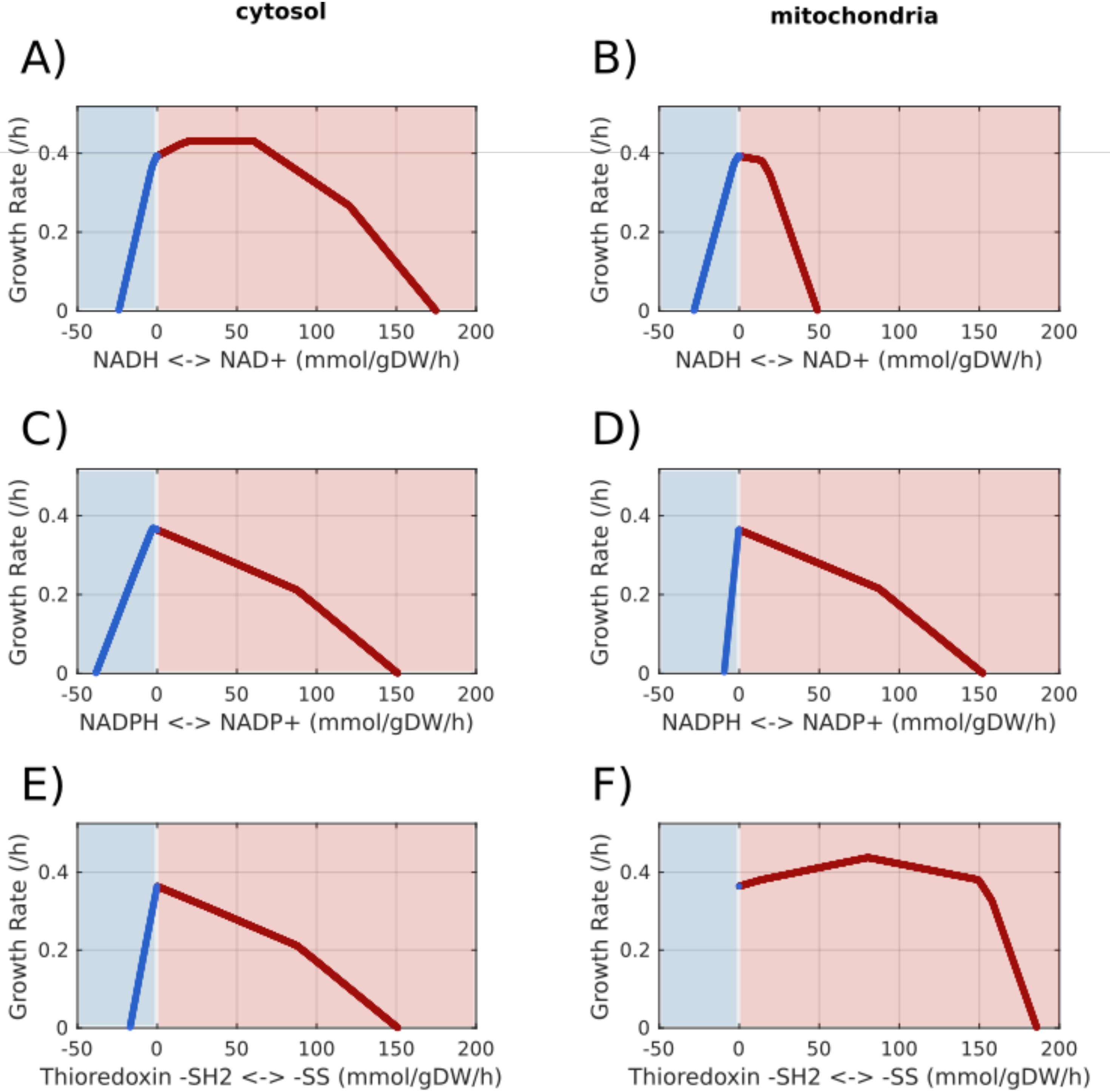

### Supplemental Figure 1

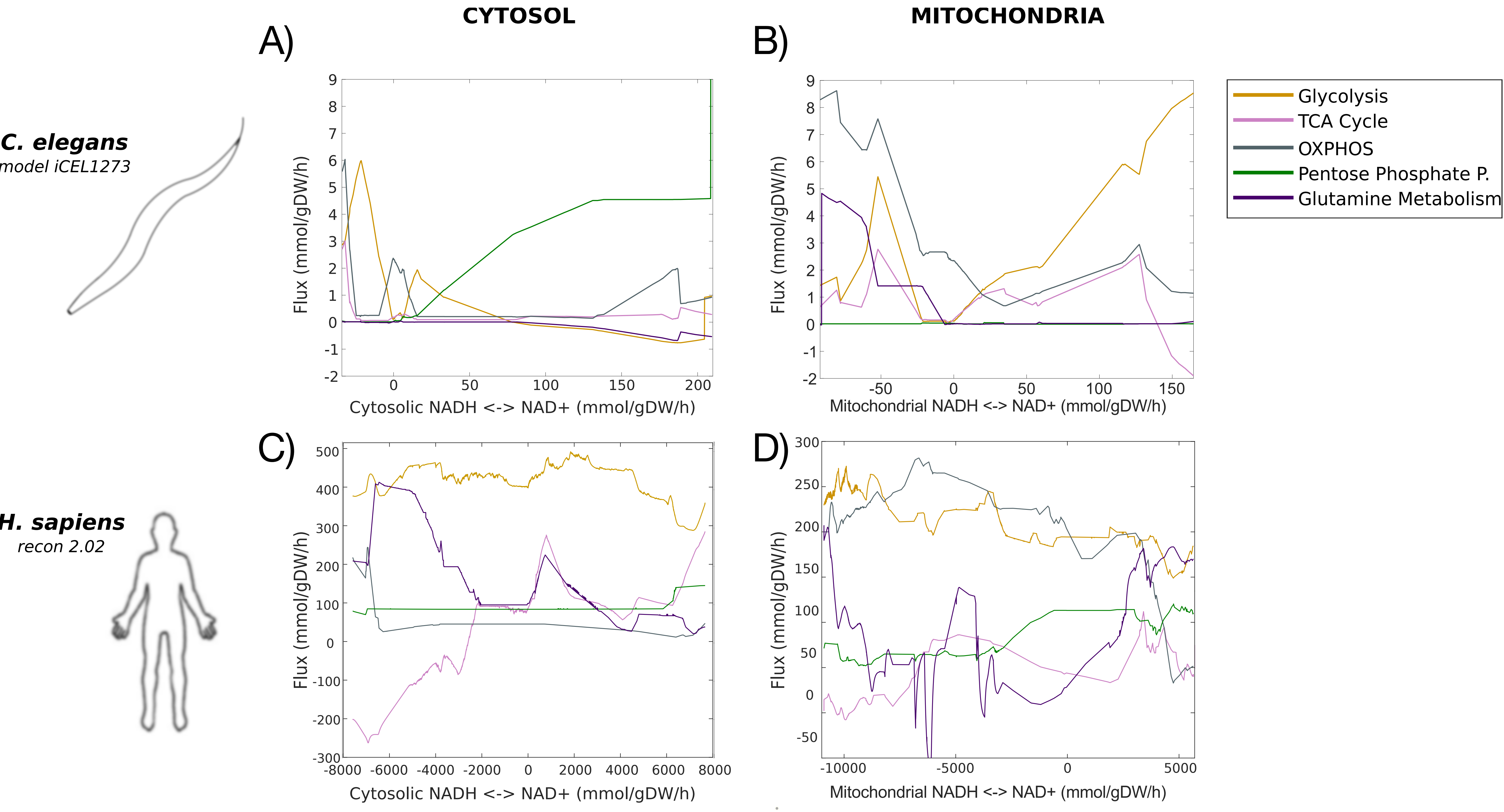
